# Inhibition of granulocyte ROS production by opioids prevents regeneration

**DOI:** 10.1101/182584

**Authors:** Elodie Labit, Lise Rabiller, Christophe Guissard, Mireille Andre, Christine Rampon, Corinne Barreau, Béatrice Cousin, Audrey Carriere, Margaux Raffin, Gilles Mithieux, Mohamad Ala Eddine, Bernard Pipy, Anne Lorsignol, Sophie Vriz, Cecile Dromard, Louis Casteilla

**Affiliations:** UMR STROMALab; Université de Toulouse, CNRS ERL5311, EFS, INP-ENVT, Inserm U1031, UPS, BP 84225 - F-31432 Toulouse Cedex 4 - France; Université Paris Diderot, Sorbonne Paris Cité Biology Department - 75205 Paris Cedex 13 – France; Centre Interdisciplinaire de Recherche en Biologie (CIRB) CNRS UMR 7241 / INSERM U1050 / Collège de France 11, Place Marcelin Berthelot - 75231 Paris Cedex 05 – France; Inserm U1213 - Lyon F-69008 - France; Université de Lyon - Lyon F-69008 - France; Université Lyon1 - Villeurbanne F-69622 - France; UMR 152-Pharma-Dev, Université de Toulouse, 31432 Toulouse, France

**Keywords:** regeneration, opioids, reactive oxygen species, MRL, adipose tissue, granulocytes

## Abstract

Inhibition of regeneration and induction of healing are classic outcomes of tissue repair in adult mammals. Here, by using gain and loss of function experiments, we demonstrate that both endogenous and exogenous opioids prevent tissue regeneration in adults, by inhibiting the early reactive oxygen species (ROS) production occurring after lesion and required for regeneration. These effects can be overcome and regeneration induced by the use of an opioid antagonist. These results, obtained in both gold-standard adult zebrafish and a newly-developed model of regeneration in adult mammals, demonstrate that this mechanism can be considered as a general paradigm in vertebrates. In addition, we show that opioids act via signaling through peripheral mu-receptors expressed on granulocytes. This work clearly demonstrates the deleterious role of opioids on tissue regeneration through the control of ROS production in vertebrates and thus questions about opioid-based analgesia in perioperative care.

## INTRODUCTION

Wound healing or regeneration are two opposite ways of tissue repair that take place after injury. While occurring in lower vertebrates and newborn mammals, regeneration after massive resection is largely impaired in adult mammals that instead exhibit wound healing ^1^. First line of defence immediately induced after the injury, inflammation plays a crucial role in the outcome of injury. Inflammation generates a well-known cascade of immune events, among which early recruitment of myeloid cells notably granulocytes on the injured site, and release of detersive molecules, as reactive oxygen species (ROS), cytokines and matrix molecules. The beneficial effect of ROS on regeneration has been mainly described in the adult zebrafish ^2-5^, newt ^6^, planarian ^7^, gecko ^8^ and xenopus tadpole ^9^.

After injury, inflammation is also associated with the peripheral release of endogenous opioid peptides by immune cells infiltrating injured tissue or neural cells ^10^. In this context, opioids play both anti-inflammatory and analgesic roles by binding to opioid receptors on immune and neural cells ^11, 12^. Opioid analogues are therefore commonly used as exogenous agents for systematic peri-operative care procedures of pain relief ^13, 14^ and for chronic pain management, including inflammatory symptoms and lesions ^15-17^. Surprisingly the consequences of their administration on regeneration have been poorly investigated, and conflicting results have been reported in animal models with a moderate epithelium injury ^18, 19, 20, 21^.

Often investigated and considered as a therapeutic target for its key role in energy homeostasis, inguinal fat pad (IFP) is a complex tissue that displays high plasticity in adults as it can undergo phenotypic (browning) or size (expansion or reduction) modifications, depending on the metabolic context ^22, 23^. It hosts a large pool of regenerative mesenchymal stem/stromal cells widely tested for their regenerative capacities in numerous clinical trials ^24, 25^. Located just under the skin, subcutaneous IFP thus represents a relevant model to study organ plasticity in adult mammals.

We hypothesized that opioids are the key factors that direct tissue injury outcome towards regeneration or healing through their control of ROS production by immune cells. To test this hypothesis, we developed gain and loss of function experiments in MRL mice well-known for their regenerative capabilities ^26^ and in non-regenerative C57BL/6 mice. In a newly-developed model of tissue lesion, relying on massive resection of IFP, we show here that following injury, opioids prevent regeneration through inhibition of ROS production. This mechanism also occurs in the caudal fin of zebrafish suggesting that it can be considered as a general paradigm in vertebrates. Opioids effects are mediated through their specific binding to mu receptors on granulocytes. Altogether our results provide a new mechanism for the inhibition of regeneration in adults, and question about the use of opioids in post-operative care.

## RESULTS

### Massive resection of IFP induces tissue regeneration or healing in MRL and C57BL/6 adult mice respectively

To investigate the cellular processes occurring in regeneration vs healing in mammals, we developed a robust and quantifiable model of tissue regeneration or healing, relying on the massive resection (around 35% of the whole tissue) of the inguinal fat pad (IFP) in adult mice. Using the specific anatomy of the IFP, the resection was systematically performed adjacent to the lymph node used as a visual reference allowing the reproducibility of the resection (Figure supplement 1A). Macroscopic and microscopic observations as well as IFP weight quantification were performed 8 weeks after surgery. As expected, spontaneous macroscopic regeneration was observed in MRL mice in contrast to C57bl/6 mice that exhibit healing (Figure 1A). Regenerative IFP exhibited adipocytes, blood vessels and nerves organized in typical shape and structure similar to the ones observed in the contralateral IFP used as an internal control. In contrast, healing IFP was characterized by the absence of adipocytes and high collagen deposition (fibrosis) revealed by second harmonic imaging (Figure 1B). Regeneration was then quantified by the regeneration index (RI) i.e. the weight ratio between the resected IFP and the uninjured contralateral IFP, having previously controlled that uninjured IFP weight did not change over the same time following unilateral IFP resection (Figure supplement 1B). Consistent with macroscopic and microscopic observations, 8 weeks after resection, RI was significantly higher in regenerative IFP than in healing IFP (0.94±0.037 *vs* 0.69±0.017 respectively) (Figure 1C). Same results were obtained as soon as 2 and 4 weeks after resection (Figure supplement 1C) and no further regeneration was observed in C57Bl/6 mice, even one year after resection (data not shown). According to these results, this newly-developed tissue lesion can be used to decipher regeneration and healing regulatory processes.

**Figure 1.**
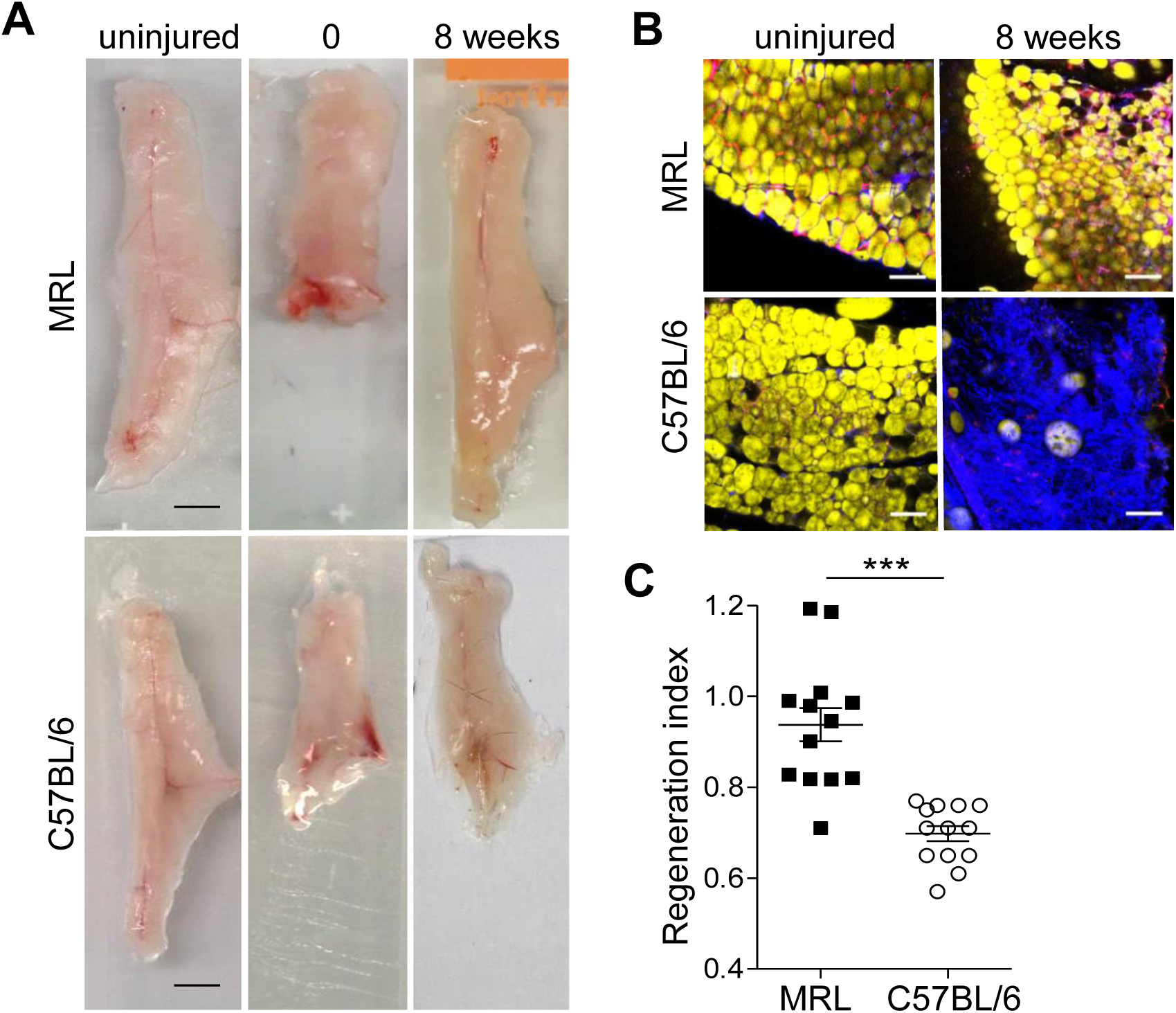
MRL but not C57BL/6 mice can regenerate inguinal fat pad (IFP). (**A**) Macroscopic view of uninjured IFP and IFP at 0 and 8 weeks post-resection in MRL and C57BL/6 mice. Scale bars: 0.5 cm. (**B**) Imaging of uninjured IFP and IFP 8 weeks post-22 resection, showing adipocytes (BODIPY staining, yellow), vascularization (lectin staining, red) and collagen deposition (second harmonic generation, blue) in MRL and C57BL/6 mice. Scale bars: 100 µm. (**C**) Quantification of IFP regeneration in MRL (black symbols) and C57BL/6 mice (white symbols) using the weight ratio (regeneration index) between the resected and the uninjured contralateral IFP 8 weeks post-resection. Data are represented as mean ± SEM. (*** p < 0.0001).

### Opioid signaling controls IFP regeneration

To investigate whether opioids inhibit regeneration and/or induce healing, spontaneous regenerative MRL mice and non-regenerative C57Bl/6 mice were treated with an opioid receptor agonist (tramadol, TRAM), or antagonist (naloxone methiodide, NAL-M) respectively. TRAM treatment induced a significant decrease in RI 4 weeks after resection (Figure 2A; 0.79±0.02 *vs.* 0.65±0.030, without vs with TRAM respectively). In contrast, RI was significantly higher in NAL-M treated mice compared to untreated mice as soon as 2 weeks after resection (Figure 2B; 0.67±0.007 *vs* 0.75±0.024 in untreated vs treated mice respectively), and this effect was amplified after 4 weeks (RI 0.69±0.013 *vs* 0.84±0.009, without vs with NAL-M treatment respectively) (Figure supplement 2A, 2B). Altogether, these results demonstrate that exogenous and endogenous opioids inhibit spontaneous tissue regeneration and favour tissue healing. Since NAL-M does not cross the blood-brain barrier, our results suggest that the anti-regenerative effect of opioids were mediated through peripheral opioid receptors.

**Figure 2.**
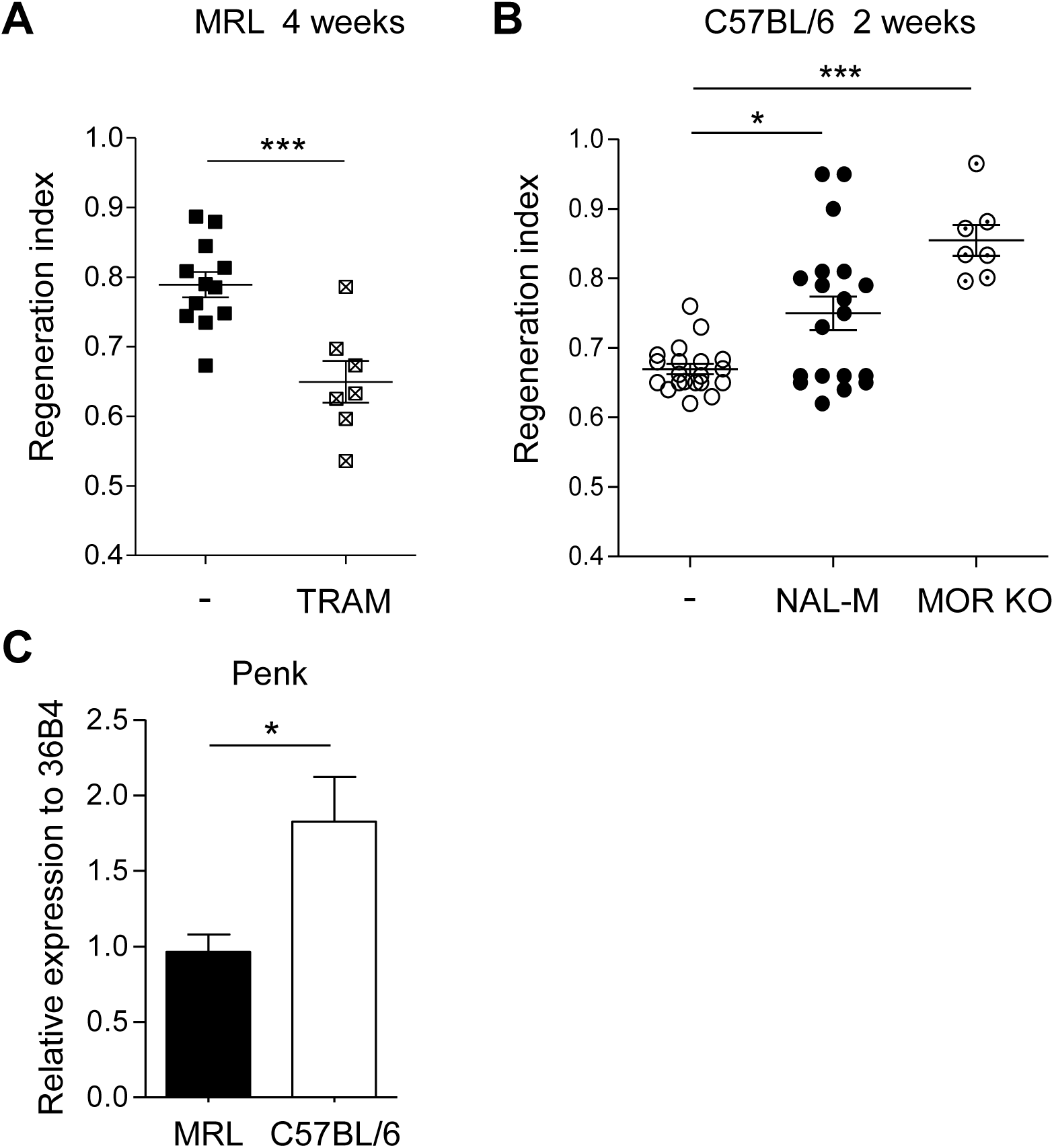
Opioid signaling controls tissue regeneration. (**A**) Quantification of IFP regeneration 4 weeks post-resection in MRL mice without (black symbols) or with (crossed symbols) treatment with the opioid receptor agonist TRAM. (**B**) Quantification of IFP regeneration 2 weeks post-resection in C57BL/6 mice without (white symbols) or with (black symbols) treatment with the opioid receptor antagonist NAL-M or in mu-opioid receptor knockout mice (MOR KO) (dotted symbols). (**C**) Penk mRNA expression in MRL (black) and C57BL/6 (white) mice IFP. Data are represented as mean ± SEM. (*p<0.05, ***p < 0.0005). TRAM: tramadol, NAL-M: naloxone methiodide.

Considering the implication of mu-opioid receptor (MOR) isoforms in the anti-inflammatory effect of opioids ^27, 28^, the involvement of MOR in regeneration was examined using MOR-deficient mice (MOR-KO). 2 weeks after resection, RI of MORKO mice was significantly higher compared to untreated C57BL/6 mice (0.85±0.022 *vs* 0.67±0.007 in MOR-KO vs untreated C57BL/6 mice respectively; Figure 2B) indicating that IFP regeneration occurred in the absence of MOR. Treatment of MORKO mice with NAL-M had no additional effect on regeneration (data not shown), suggesting that NAL-M exerts its effects only through MOR.

We thus postulated that the absence of spontaneous regeneration in C57BL/6 compared to MRL mice could be associated with a higher tonus of endogenous opioids in the IFP of C57BL/6 mice. Consistent with this hypothesis, the IFP-expression of proenkephalin (PENK, enkephalin opioid peptide precursor) was significantly higher in IFP of C56Bl/6 than in MRL mice (Figure 2C). In contrast, neither prodynorphyn (PDYN, dynorphine opioid peptide precursor) nor pro-opiomelanocortin (POMC, endorphine opioid peptide precursor) mRNA were detected in IFP in both mice strains (data not shown). Altogether these data demonstrate that opioids inhibit regeneration in adult mammals through the peripheral MOR. This effect could be mediated by endogenous PENK.

### Opioid signaling prevents regeneration through control of ROS levels in zebrafish

To demonstrate that the anti-regenerative effects of opioids were not tissue and/or model dependent, we used the gold-standard adult zebrafish caudal fin regeneration model ^29^. Fish were incubated in NAL-M or TRAM from the time of amputation to analysis, and the size of the regenerate was quantified after amputation (Figure 3A). As in our mice model, NAL-M enhanced the regenerate size (Figure 3B, 3C, 137.7%±26), while TRAM inhibited regeneration (Figure 3B, 3C, 70±13%). These results demonstrate that inhibitory effect of exogenous and endogenous opioids on tissue regeneration is a prevailing process in adult vertebrates.

**Figure 3.**
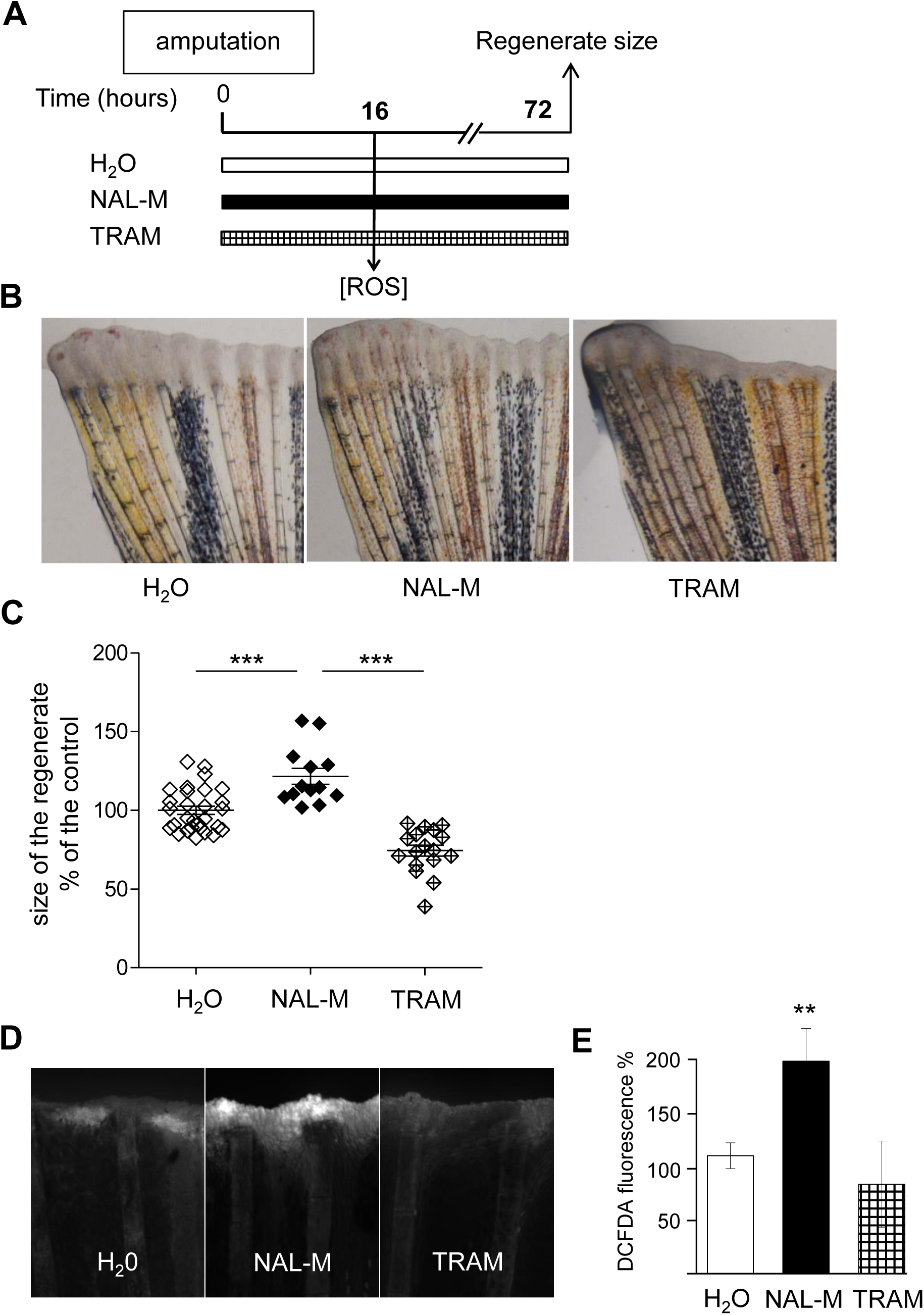
Opioid signaling prevents regeneration through control of ROS production in zebrafish. (**A)** Scheme of the experiment. Caudal fins of adult fish were amputated and then allowed to regenerate for 16 or 72 hours. (**B**) Representative images 72 hpa (hours post-amputation) of caudal fins challenged to regenerate in the presence of an opioid receptors antagonist (NAL-M) or agonist (TRAM). (**C**) Quantification of the size of the regenerated tissue at 72 hpa in the control (H20) (white symbols) and in fish treated with NAL-M (black symbols) or TRAM (crossed symbols). (**D**) ROS detection (representative images) at the level of the amputation plane at 16 hpa. (**E**) ROS quantification. Data are represented as mean ± SEM. (**p < 0.01; ***p < 0.0001). NAL-M: naloxone methiodide, TRAM: tramadol.

In zebrafish, regeneration has been widely demonstrated to be controlled by ROS ^3^. We thus postulated that opioids control caudal fin regeneration through regulation of ROS production. NAL-M treatment enhanced ROS levels at the tip of the amputated fin, the major site of ROS production after amputation (Figure 3D, 3E) while TRAM reduced the overall ROS production, even when the area of ROS detection was extended compared to the control (Figure 3D, 3E). These results obtained in the caudal fin of zebrafish demonstrate that opioids inhibit tissue regeneration via abolishing the transient ROS peak.

### Opioid signaling prevents regeneration through control of ROS levels in adult mammals

We therefore aimed to demonstrate that regeneration in adult mammals was associated with ROS production. In line with this hypothesis, IFP resection induced a robust and transient peak of ROS in the injured tissue of regenerative MRL mice and not C57Bl/6 mice as shown by *in vivo* imaging using luminol (Figure 4A). ROS production in MRL mice reached maximal values at 12 hours after resection and returned to normal values 72 hours following surgery (Figure 4B). MRL mice treatment with apocynin (APO, inhibitor of NADPH p47 phox subunit translocation ^30^ after IFP resection, induced a severe decrease in RI 2 weeks after surgery (Figure 4C; 0.85±0.027 vs. 0.62±0.019 in untreated vs APO treated IFP). These results demonstrate that a robust and transient peak of ROS is required for proper tissue regeneration in adult MRL mice.

**Figure 4.**
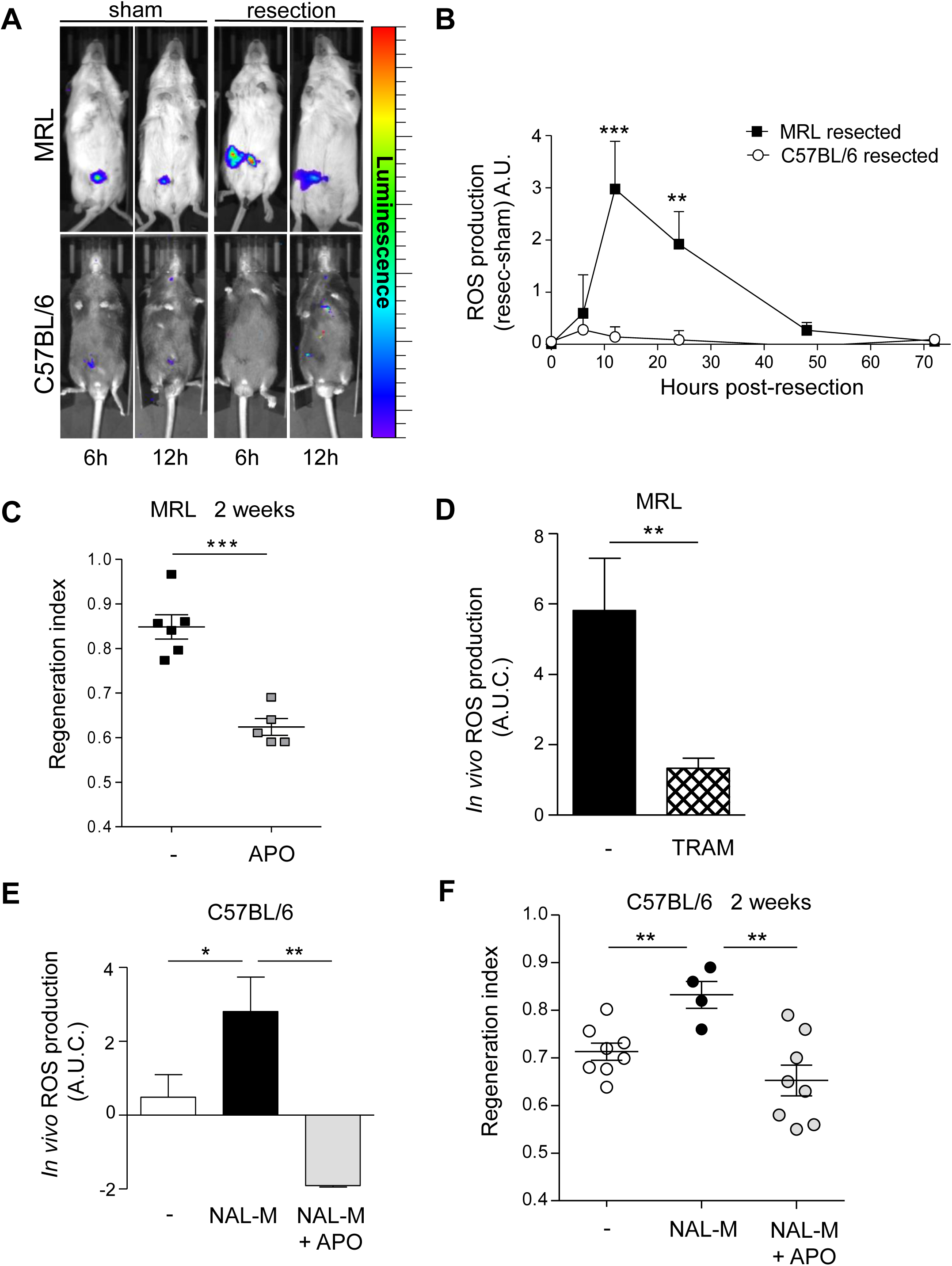
ROS production is required for IFP regeneration. (**A**) Representative *in vivo* imaging of ROS production at 6 and 12 hours after surgery in sham and resected MRL and C57BL/6 mice. (**B**) *In vivo* quantification of ROS production at 0, 6, 12, 24, 48 and 72 hours post-resection in MRL (black symbols) and C57BL/6 (white symbols) mice. n = 8 per group. A.U.: arbitrary units. (**C**) Quantification of IFP regeneration in MRL mice 2 weeks post-resection without (black symbols) or with (grey symbols) treatment with APO. Data are represented as mean ± SEM. (**p < 0.01, *** p < 0.0001). (**D**) Quantification of ROS production *in vivo* from 0 to 96 hours post-resection in MRL mice treated with TRAM. n = 8 per group. (**E**) Quantification of ROS production *in vivo* from 0 to 96 hours post-resection in C57BL/6 mice without (white bar) or with (black bar) NAL-M treatment or with NAL-M and APO treatment (grey bar). n = 8 per group. (**F**) Quantification of IFP regeneration 2 weeks post-resection in C57BL/6 mice without (white symbols) or with (black symbols) NAL-M treatment or with NAL-M and APO treatment (grey symbols). Data are represented as mean ± SEM. (* p < 0.05, **p<0.005, ***p < 0.0005). APO: apocynin. TRAM: tramadol, NAL-M: naloxone methiodide.

The effects of opioids on ROS production were then investigated *in vivo* by treatments with antagonists or agonists of opioid receptors. *In vivo* imaging revealed that the ROS peak observed after resection was strongly inhibited in IFP of MRL mice within 96 hours after TRAM treatment (Figure 4D), and this was associated with a reduced RI (Figure 2A). In contrast, NAL-M treatment of C57Bl/6 mice induced a significant increase in ROS production and subsequent increase in RI, 2 weeks post-resection (Figure 4E, 4F). This opioid effect on ROS production was abolished after APO treatment, leading to a severe decrease in RI (Figure 4E, 4F).

Taken together, these data demonstrate that ROS production is required for proper tissue regeneration in adult mammals and that opioids inhibit this regeneration through the control of ROS production.

### Mu-opioid receptors on granulocytes mediate opioids anti-regenerative effect

Several evidences suggest that immune cells may be necessary to induce regeneration ^31-33^. Immune cells were therefore quantified in the IFP, in both MRL and C57Bl/6 mice after resection. Flow cytometry analysis of the IFP area located close to the wound showed that the number of granulocytes (CD45^+^/Gr-1^+^) was significantly increased in NAL-M treated mice compared to untreated mice whereas no change in macrophages (CD45^+^/CD11b^+^/F4/80^+^) or mast cell (CD45^+^/CD11b-/CD117^+^/FcεRI^+^) populations was detected (Figure 5A). This indicates that IFP regeneration correlates with accumulation of granulocytes. Specific depletion of this cell population was thus undertaken before resection and NAL-M treatment in C57BL/6 mice. Injection of an anti Gr-1 blocking antibody induced a significant and specific decrease in Gr-1^+^ cell number in IFP within 24 hours (Figure supplement 3A) whereas no significant change in mast cells and macrophages numbers was observed. In the absence of granulocytes, the ROS production after IFP resection and NAL-M-treatment was severely decreased (Figure 5B) as well as the RI (0.61±0.03) (Figure 5C). This suggests that granulocytes play a key role in the control of regeneration via the production of ROS.

**Figure 5.**
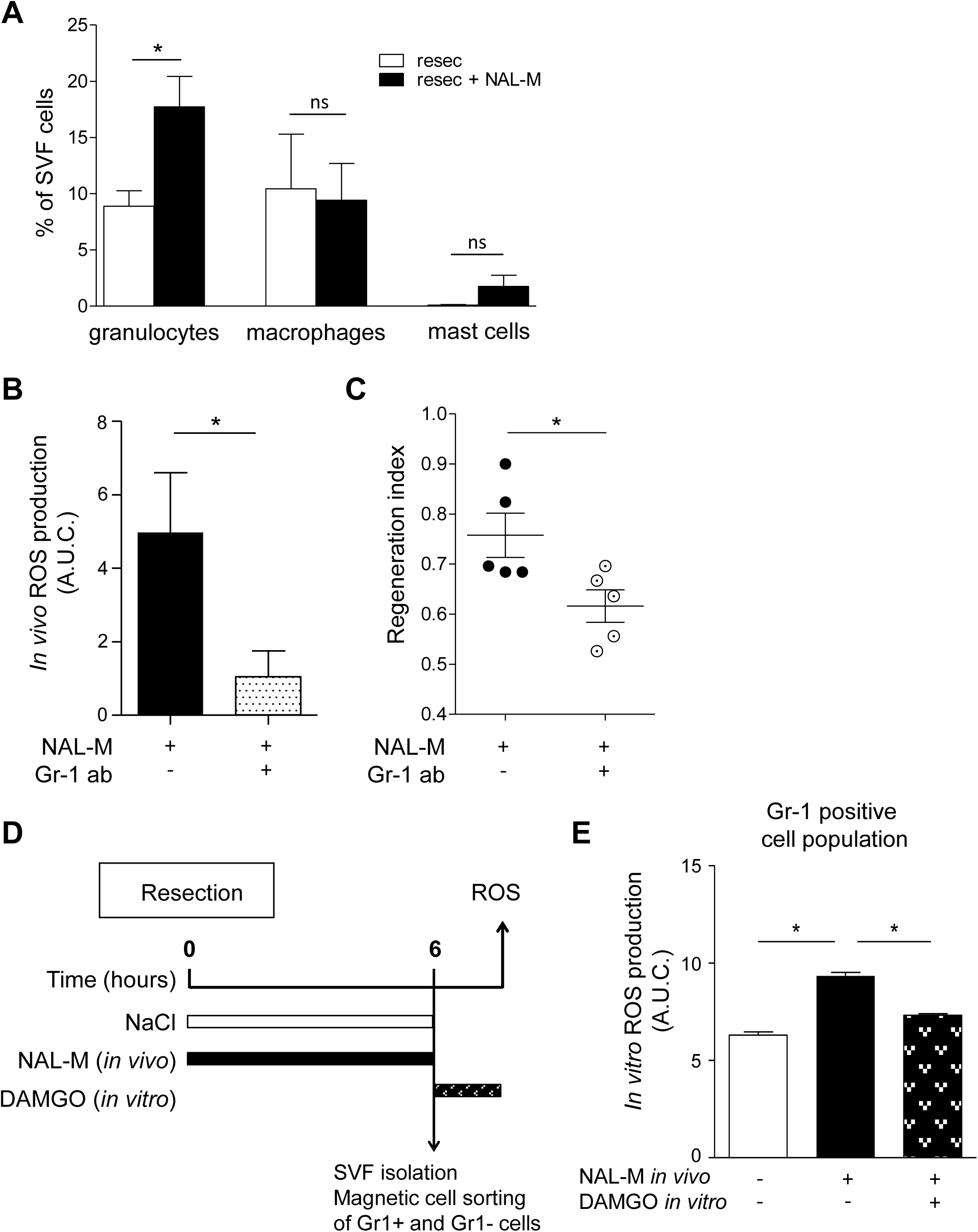
Mu opioid receptors on granulocytes mediate anti-regenerative effect of opioids on IFP. (**A**) Granulocytes (CD45^+^Gr-1^+^), macrophages (CD45^+^CD11b^+^F4/80^+^) and mast cells (CD45^+^CD11b^-^CD117^+^FcεRI^+^) populations in the stromal vascular fraction of IFP of C57BL/6 mice, 6 hours post-resection without (white bars) or with (black bar) treatment with NAL-M. (**B**) Quantification of *in vivo* ROS production from 0 to 72 hours post-resection in C57BL/6 mice treated with NAL-M, without (black bar) or with (dotted bar) Gr-1 blocking antibody. n = 5 per group. (**C**) Quantification of regenerated IFP 4 weeks post-resection of C57BL/6 mice treated with NAL-M, without (black symbols) or with (dotted symbols) Gr-1 blocking antibody. (**D**) Scheme of the experiment. C57BL/6 mice were treated with NAL-M or NaCl immediately post-resection of IFP and the Gr-1 positive and Gr-1 negative populations were sorted 6 24 hours post-resection. ROS production was measured in sorted cells treated *in vitro* with the mu-opioid receptors selective agonist DAMGO. (**E**) *In vitro* quantification of ROS production in the Gr-1 positive cell population isolated from resected IFP. n = 3 per group. Data are represented as mean ± SEM. (* p < 0.05). SVF: stromal vascular fraction, NAL-M: naloxone-methiodide, DAMGO: [D-Ala^2^, N-MePhe^4^, Gly-ol]-enkephalin.

Finally, to determine whether opioids exert a direct effect on granulocytes, Gr1^+^ and Gr1^-^ cells were sorted from C57Bl/6 mice 6 hours after IFP resection with or without NAL-M treatment (Figure 5D). *In vitro* ROS production was quantified by using luminol. Gr-1^+^ cell sorted from NAL-M treated mice exhibited a significant higher ROS production than Gr-1^+^ cells sorted from untreated mice (Figure 5E), showing that IFP regeneration correlates with ROS production by granulocytes. *In vitro* treatment of Gr-1^+^ cells with the mu-opioid receptors selective agonist DAMGO abolished the NAL-M induced ROS production (Figure 5E). No variation in ROS production was measured in Gr-1^-^ sorted cells (Figure supplement 3B). Altogether, these data clearly demonstrate a direct inhibitory effect of opioids on ROS production by Gr1^+^ cells after IFP resection, leading to inhibition of IFP regeneration.

## DISCUSSION

By using convergent and complementary models, and loss and gain of function approaches, we show for the first time in adult mammals, that in the early steps post-injury opioids prevent regeneration processes via the inhibition of an early burst of ROS produced by granulocytes, through a signaling via mu-receptors.

A transient but early use of an opioid receptor agonist is sufficient to inhibit spontaneous regeneration of the IFP in the regenerative strain of mice MRL, when treatment with an antagonist is able to induce regeneration in the non-regenerative strain of mice C57BL/6. This is in line with a very recent study suggesting that administration of morphine (opioid receptor agonist) delays pancreatic regeneration after acute pancreatitis in mice ^34^. However, our data demonstrates also that endogenous opioids are key determinant in regeneration control. Indeed, endogenous PENK expression is lower in regenerative MRL vs non regenerative C57Bl/6 mice. In addition, blocking endogenous opioid receptors (without any administration of exogenous opioids) is sufficient to induce tissue regeneration.

We show here that a strong production of ROS by NADPH oxidation occurs both in the spontaneous and the pharmacologically-induced regeneration processes and that its inhibition systematically prevents tissue regeneration in adult mice. We show that this strong production of ROS is a very early and transient phenomenon. This is the first demonstration of the crucial role of ROS in the control of regeneration in adult mammals. In agreement with our data, a correlation between ROS production and regeneration was described in adult acomys mice ^33^. In addition, several reports have highlighted the importance of ROS in regeneration in invertebrates and lower vertebrates ^3, 9, 35^. Altogether, this shows that ROS requirement for proper regeneration is a general paradigm in vertebrates including adult mammals.

Our findings support an anti-oxidative effect (inhibition of ROS production) of endogenous opioids *in vivo* after tissue lesion that prevents regeneration. *In vivo* such a role of opioids has never been described. An opposite effect of long term opioid treatment was however described in cultured cells (for review see ^36^).Interestingly, our data demonstrate the inhibitory role of opioids on regeneration, through the inhibition of ROS production in different tissues (IFP and caudal fin) and animal models (mice and zebrafish), showing that this mechanism is not tissue or model specific, and can be considered as a general paradigm in vertebrates.

Our data undoubtedly show that granulocytes play a key role in the very early steps of regeneration in mammals. Granulocytes, notably neutrophils are among the first immune cells to infiltrate injured tissue. Their role in regeneration has been suggested in zebrafish ^37^, but remains controversial. They have also been shown to be the principal regulators of optic nerve regeneration in mice ^38^. In view of the increased number of granulocytes at the site of lesion after NAL-M treatment, we can speculate that endogenous opioids through activation of the Gi-protein coupled MOR prevent chemokine secretion required for granulocytes recruitment as it was described after morphine treatment ^27^. Although granulocytes are the only cells to accumulate in the first hours after IFP resection, it does not rule out a possible role for other immune cells in the whole process. Indeed, macrophages have been shown to be involved in regeneration in mammals, at least 48 hours after lesion ^32, 33^ suggesting that macrophages are enrolled at later stage of regeneration process.

In conclusion, we demonstrate a pivotal role of opioids in the tissue repair outcome after lesion in adult vertebrates. Both endogenous and exogenous opioids inhibit regeneration very early in the process *via* inhibition of granulocytes ROS generation. These effects can be reversed using opioid receptor antagonists. Considering that all peripheral tissues potentially contain endogenous opioids after lesion and their receptors, we propose that these results could have broad implication for mammalian tissue repair and regeneration. Finally, opioids are largely used in peri-operative care in human, and their potential role on tissue repair outcome has never been questioned. Our results open therefore new perspectives to develop new and pro-regenerative perioperative care protocols that could enhance tissue regeneration while preventing pain and inflammation.

## MATERIALS AND METHODS

### Animals

All experiments in mice were performed on 5- to 7-weeks-old male mice. C57BL/6 mice were obtained from Harlan Laboratories. MRL/MpJ mice were obtained from The Jackson Laboratory and bred in the CREFRE-US006 (Centre Régional d’Exploration Fonctionnelle et Ressources Expérimentales). Congenic male C57BL/6J CD45.1 mice were provided from ENVIGO. MOR-KO mice were a gift from G. Mithieux. Animals were group-housed (3 or 4 per cage) in a controlled environment (12-hour light/dark cycles at 21°C) with unrestricted access to water and a standard chow diet in a pathogen-free animal facility (IFR150). Animals were maintained in accordance to guidelines of the European Community Council. Mice were killed by cervical dislocation. All experiments were carried out in compliance with European Community Guidelines (2010/63/UE) and approved by the French ethics committee (protocol reference: CEEA-122 2014-66).

Zebrafish colonies (AB-Tu) were maintained using standard methods. The animal facility obtained approval from the French agreement from the Ministère de l’agriculture (n° C75-05-12), and the protocols were approved by the Ministère de l’éducation nationale de l’enseignement supérieur et de la recherche (00477.02). To maintain a healthy colony, a cycle of 14 h light-10 h dark was used, and a water temperature of 28°C was maintained, with a maximal density of five fish per liter. Water filtration depended on Aquatic Habitat stand-alone fish housing and operated automatically (Aquatic Habitat, Inc., FL, USA). Fish were fed twice per day with live 2-day-old artemia. For manipulation and amputation, the adult zebrafish (5-10 months of age) were anesthetized in 0.1% Tricaine (ethyl-m-aminobenzoate), the caudal fins were amputated at the level of the first ray bifurcation and the fins were allowed to regenerate for various lengths of time. The efficiency of regeneration was quantified at 72 hpa. The surface of the blastema was measured and subsequently divided by the square length of the amputation plane for each fish. The efficiency of regeneration is expressed as a percentage of the control.

### IFP resection

Control mice were used for the baseline control and did not undergo surgery. Mice underwent unilateral resection of subcutaneous IFP. Animals were anesthetized by inhalation of isoflurane and intraperitoneal (i.p.) administration of ketamine (50 mg/kg, 50 µl, Virbac, Carros, France). After the mice were shaved, a single incision was made on the abdomen to access and excise 35 to 40% of the right inguinal fat pad between lymph node and groin. The left IFP was not submitted to a surgical procedure and was thus used as internal control. Sham animals were shaved and opened. For these two groups (sham and resected), the skin was closed with 3 suture points. To quantify IFP regeneration, the weight ratio between right (i.e with resection) and left (i.e. contralateral) fat pads was calculated (regeneration index).

### Immunohistochemistry

IFP sections of 300 µm thickness were incubated in blocking solution (2% Normal Horse Serum and 0.2% triton X-100 in PBS) at room temperature (RT) and incubated for 24 hours at RT with bodipy 558/568 (1:1000, D3835, Invitrogen, Carlsbad, CA, USA) and lectin biotinylated (1:100, B-1105, AbCys, Courtaboeuf, France). Then sections were incubated overnight at 4°C with Streptavidin Alexa-647 (1:100, S21374, Invitrogen, Carlsbad, CA, USA), mounted on a coverslip and imaged using a Confocal Laser Scanning microscope (LSM780, Carl Zeiss, Oberkochen, Germany). Second harmonic generation (SHG) signal allowing visualization of fibrillar collagen was acquired using a mode-locked 890 nm laser. SHG was measured at 445 nm with a 20 nm bandpass filter. Images were processed using Fiji software (NIH, Bethesda, MD, USA).

### *In vivo* treatments

Mice were treated on days 0-4 after IFP resection by subcutaneous injection of naloxone methiodide (NAL-M) (17 mg/kg, N129, Sigma Aldrich, Saint Louis, MO, USA), naloxone hydrochloride dehydrate (NAL) (17 mg/kg, N7758, Sigma Aldrich, Saint Louis, MO, USA) or Acetovanillone Apocynin (100 mg/kg, 200 µl s.c., W508454, Sigma Aldrich, Saint Louis, MO, USA). Tramadol was added to the drinking water (10 mg/kg, Grunenthal, Belgium). Granulocytes depletion was achieved by i.p. injection of anti-mouse Gr-1 blocking antibody (Gr-1 ab) (25 µl i.p., BE0075, BioXCell, West Lebanon, NH, USA) 24 hours before IFP resection. Fish were incubated in naloxone methiodide (NAL-M) (5 μM, N129, Sigma Aldrich, Saint Louis, MO, USA) or Tramadol (1 mM, Grunenthal, Belgium).

### *In vivo* ROS imaging

Mice were briefly anesthetized by inhalation of isoflurane and i.p. injected with 5 mg of luminol (5-amnio-2,3-dihydro-1,4-Phtalazinedione, A4685, Sigma Aldrich, Saint Louis, MO, USA) in 100 µl of PBS. Animals bioluminescence was imaged using an IVIS Spectrum 200 (Caliper Life Science, Hopkinton, MA, USA) for 2 min exposures at different times after luminol injection. Image analyses were performed using LivingImage 3.0 Software (Caliper Life Science, Hopkinton, MA, USA). The images were calibrated with intensity colour from 30 (min) to 330 (max). For each animal, the sham surgery area signal was subtracted to resected area photon flux.

In zebrafish, 2’,7’-dichlorodihydrofluorescein diacetate (H2DCFDA, Calbiochem, San Diego, CA, USA) was used to monitor the accumulation of ROS in adult zebrafish fins. Fluorescent DCF was formed through ROS oxidation. Zebrafish were incubated with H2DCFDA (50 µM) 2 h prior to confocal imaging. Spinning-disk images were acquired using a 4x/0.15 N.A. objective on a Nikon Eclipse Ti microscope equipped with a CoolSnap HQ2/CCD camera (Princeton Instruments, Trenton, NJ, USA) and a CSUX1-A1 (Yokogawa) confocal scanner. MetaMorph software (Molecular Devices, Sunnyvale, CA, USA) was used to collect the data. Fluorescence was excited with a 491 nm laser and detected with a 525/39 nm filter. Quantification of fluorescence intensity was performed using ImageJ software.

### Isolation of adipose derived stromal vascular cell fraction

Inguinal fat pad from C57BL/6 mice was carefully dissected, mechanically dissociated and digested at 37°C with collagenase (Roche Diagnostics, Mannheim, Germany) for 30 minutes. After elimination of undigested fragments by filtration (25 µm), cells were collected by centrifugation at 1800 rpm for 10 minutes. Cells obtained from both tissues were incubated for 5 minutes in hemolysis buffer (140 mM NH_4_Cl and 20 mM Tris, pH 7.6) to eliminate red blood cells and washed by centrifugation at 1600 rpm for 6 min in PBS. The cells were counted and used for flow cytometric analysis or sorting before plating *in vitro*.

### Cell phenotyping and sorting

Flow cytometry was used i) to quantify macrophages, mast cells and granulocytes following IFP resection and ii) to check efficiency of the granulocytes depletion in experiments with Gr1 antibody injection. Freshly isolated cells from the stroma-vascular fraction (SVF) were stained in PBS containing FcR-blocking reagent. Phenotyping was performed by immunostaining with conjugated CD45 APC-H7 (Clone104), CD11b-PE-Cy7 (clone M1/70), Gr-1-APC (Ly6G/Ly6C) (RB6-8C5), F4/80-PerCP Cy5,5 (Clone BM8), c-Kit-APC (clone 2B8) and FcεRIα PE (Clone MAR-1) and compared with isotype-matched control mAb (BD Biosciences). Cells were washed in PBS and analyzed on a LSR Fortessa flow cytometer (BD Biosciences). Data were acquired with FACSDiva Version 7 software (BD Biosciences) and analyzed using Kalusa Version 1.3 software (BeckmannCoulter). For granulocyte cells isolation, magnetic sorting was used: 6 hours after IFP resection, SVF from adipose tissue was prepared; cells were incubated with 10 µl of anti-Gr-1 microbeads (Miltenyi Biotec, 130-092-332) and sorted with MACSQuant Tyto (Miltenyi biotec). Gr1^+^ and Gr1^-^ cells were maintained in HBSS during ROS level quantification by luminometer.

### *In vitro* ROS quantification

The oxygen-dependent respiratory production of 140 000 AT Gr1^-^ or Gr1^+^ sorted cells was measured by chemiluminescence in the presence of luminol (66 µM, Sigma-Aldrich) using a thermostatically monitored luminometer (37°C) (210410A EnVision Multilabel Reader). DAMGO ([D-Ala^2^, N-Me-Phe^4^, Gly^5^-ol]-Enkephalin acetate salt, E7384, Sigma Aldrich) was added 5 minutes before luminescence measure. The luminol detects both reactive oxygen and nitrogen intermediates (O_2.^-^_, ONOO^-^, OH.). Chemiluminescence was continuously monitored for 1 hour. The ROS level was quantified and compared between different conditions using the area under the curve.

### RNA Extraction and Real-Time PCR

For mouse tissues, total RNA was isolated by Qiazol extraction and purification was done using RNeasy microcolumns (Qiagen). 250 ng of total RNA were reverse-transcribed using the High Capacity cDNA Reverse Transcription kit (Life Technologies/Applied Biosystem), SYBR Green PCR Master Mix (Life Technologies/Applied Biosystem), and 300 nmol/L primers on an Applied Biosystem StepOne instrument. Penk relative gene expression was determined using the ΔΔCT method and normalized to *36B4* level.

**Table 1.**
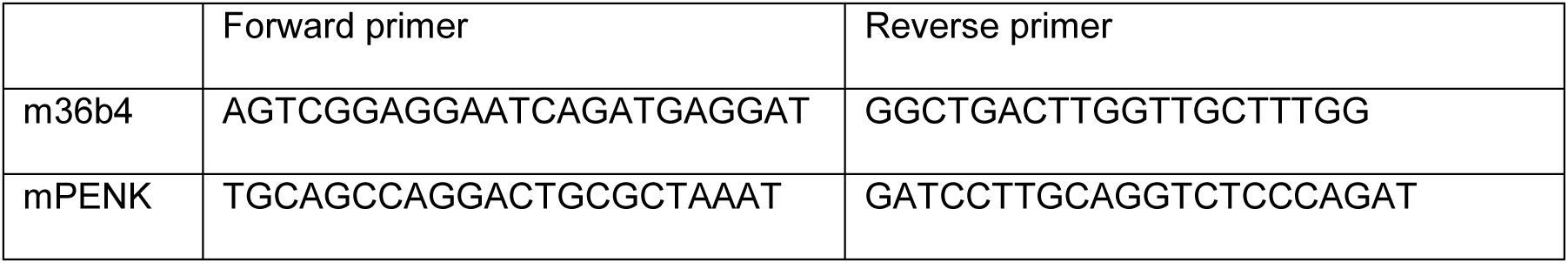
Primers sequences.

### Statistical analyses

Studies were not randomized and analyses were blinded to investigators. All results are given as means +/-SEM. Data were analyzed using an ANOVA test (when more than 2 groups) and variance between groups was compared. Statistical differences were measured using an unpaired two-sided Student’s-t-test, or a nonparametric test (Mann-Whitney) when variances between groups were different. All statistical analyses were performed in GraphPad Prism 5.0 software and a two-tailed P value with 95% confidence interval was acquired.

## Author contributions

E.L. designed the experimental approach, performed experiments, analyzed data and drafted the manuscript. L.R. performed experimental work on ROS and granulocytes in MRL and C57BL/6 mice. C.G. set up the IFP model. M.A. performed experiments on ROS and granulocytes. C.R. performed experiments on zebrafish. C.B. carried out tissue imaging. B.C. designed the experimental approach for immune cells repopulation and analyzed data. A.C. analyzed data and developed the hypothesis. M.R. and G.M. designed and analyzed data for the MOR-KO mice. M.A. performed experiments on ROS. B.P. designed the experimental approach on ROS and analyzed data. A.L. designed the experimental approach, analyzed data and drafted the manuscript. S.V. designed experiments on zebrafish, analyzed data and drafted the manuscript. C.D. designed the experimental approach, performed experiments, analyzed data and drafted the manuscript. L.C. coordinated the 19 experiments, designed the experimental approach, analyzed data and drafted the manuscript.

## Acknowledgements

The authors thank i) the US006/CREFRE INSERM/UPS (Toulouse, France), specifically the zootechnical core facility for animal care and chirurgical investigation, and ii) the Institut des Techniques Avancees du Vivant (ITAV, USR3505) for tissue imaging. The authors are grateful to B. Segui for Gr-1 blocking antibody gift and for insightful discussions, X. Sudre, S. Gandarillas and Y. Barreira for animal cares, H. Lulka for help with luminomer, C. De Vecchi for technical assistance, J. Rouquette for imaging assistance, N. Espagnolle and A. Varin for help in cell sorting and C. Sengenes for insightful discussions. This work was financially supported through grants from the EU FP7 project DIABAT (Health-F2-2011-278373), the Bettencourt Schueller Foundation and the Midi-Pyrénées region (DESR/12052900 and DESR/14050455). E. Labit and L. Rabiller received a fellowship from the French Ministère de l’Enseignement et de la Recherche.

